# ANTIBACTERIAL ACTIVITY OF CHINESE CABBAGE (Brassica rapa subsp. chinensis) AGAINST SELECTED FOODBORNE PATHOGENS

**DOI:** 10.1101/2025.08.04.668494

**Authors:** Von Jay Maico G. Gabucan, Concepcion Kristy C. Revilla, Stephanie R. Quinoviva, Dynah Marie S. Armada, Angelica Roge S. Ilog, Ana Lou Grace S. Manluyang, Ferlien Mae B. Brieta

## Abstract

Foodborne pathogens pose significant global health risks, contributing to foodborne illnesses and antimicrobial resistance. The increasing demand for plant-derived alternatives to synthetic antimicrobials highlights the relevance of investigating locally available vegetables with potential antibacterial properties. This study aimed to evaluate the antibacterial activity of the ethanolic extract of Chinese cabbage (*Brassica rapa* subsp. *chinensis*) against selected foodborne pathogens and to identify the functional groups present in the extract using Fourier-Transform Infrared (FTIR) spectroscopy. Fresh Chinese cabbage leaves were dried, powdered, and extracted using 80% ethanol. The antibacterial activity was assessed through the Kirby-Bauer disk diffusion method against *E. coli, S. enteritidis*, and *S. aureus*, with ciprofloxacin and vancomycin as positive controls. FTIR analysis revealed the presence of alcohol, alkene, amine, and alkyl functional groups, indicating the presence of potentially bioactive phytochemicals. In the antibacterial assay, the extract produced mean zones of inhibition of 24.18⍰±⍰0.305 mm for *S. enteritidis*, 9.49⍰±⍰2.726 mm for *S. aureus*, and 7.78⍰±⍰2.019 mm for *E. coli*. Among the tested pathogens, *S. enteritidis* exhibited the highest sensitivity to the extract. However, all inhibition zones were significantly smaller and statistically different than those of the positive control antibiotics, indicating moderate antibacterial activity. The findings support the antibacterial potential of *Brassica rapa* subsp. *chinensis*, particularly against *S. enteritidis*. The presence of functional groups associated with antimicrobial properties suggests its suitability as a natural antibacterial agent. Further phytochemical analysis and formulation studies are recommended to enhance its efficacy and explore its potential applications in food safety and alternative medicine.

## INTRODUCTION

Foodborne diseases remain a significant global public health concern. These illnesses account for nearly 33 million disability-adjusted life years (DALYs), with children under five years old disproportionately affected, comprising nearly 30% of the mortality burden (World Health Organization, 2015). The global burden of foodborne diseases has been found to be comparable to that of major infectious diseases such as HIV/AIDS, malaria, and tuberculosis in terms of overall impact on health and productivity (Havelaar et al., 2015). In the United States, the Centers for Disease Control and Prevention (CDC) estimate that foodborne pathogens cause about 48 million illnesses, 128,000 hospitalizations, and 3,000 deaths each year (Scallan et al., 2011). The associated economic impact, including direct medical costs and productivity losses, ranges between 5 to 17 billion US dollars annually (Hoffmann et al., 2012).

In the Philippine context, food safety remains a critical public health concern, particularly in urban centers where informal street vending is widespread and regulatory oversight is limited. Street foods provide affordable nutrition for low-income populations, yet numerous studies have confirmed their vulnerability to microbial contamination due to inadequate sanitation, poor hygiene practices, and improper food storage. Street-vended chicken and pork products in Laguna were shown to contain unacceptably high levels of aerobic microbes, *E. coli, Staphylococcus aureus*, and *Salmonella*, signaling significant risk for disease transmission (Manguiat & Fang, 2013). In a news reported in Davao City, food samples collected from 120 street vendors revealed contamination with *Escherichia coli* and *Salmonella*, both of which are leading causes of foodborne illness (Bangcongco, 2012). Similar findings were reported that identified disease causing bacteria among sauces used in street foods in Cagayan de Oro City (Lubos, 2012). Moreover, food safety in the Philippines is fragmented and poorly coordinated with regulatory systems, which leads to overlapping responsibilities, enforcement gaps, and a largely reactive approach to foodborne illness outbreaks (Collado et al., 2015). These challenges underline the need for affordable, community-accessible interventions, including plant-based antimicrobials that could supplement or provide alternatives to conventional antibiotics — especially in light of the rising threat of antimicrobial resistance.

The increasing prevalence of antimicrobial resistance (AMR) among foodborne pathogens adds urgency to the search for alternative antibacterial agents. The World Health Organization has recognized AMR as a growing public health threat, necessitating the exploration of new antimicrobial sources, including plant-based compounds.

Plants from the Brassicaceae family have gained attention for their potential antimicrobial properties. These plants are rich in secondary metabolites such as glucosinolates, flavonoids, and phenolic compounds, which may exhibit antibacterial, antioxidant, and anticancer properties (Hu et al., 2004; Johnson, 2002; Sánchez-Pujante et al., 2017). When hydrolyzed, glucosinolates produce biologically active compounds such as isothiocyanates, thiocyanates, and indoles, which have demonstrated inhibitory activity against a wide range of microbial species (Kyung & Fleming, 1997). Specifically, Chinese cabbage (*Brassica rapa* subsp. *chinensis*), a readily available leafy vegetable in Southeast Asia, contains S-methyl-L-cysteine sulfoxide—a precursor to methyl methanethiosulfinate, a compound known for its antimicrobial action.

Given its accessibility and nutritional value, Chinese cabbage holds promise as a low-cost source of antibacterial agents. However, scientific data on its efficacy against common foodborne pathogens remain limited. This study, therefore, aims to evaluate the antibacterial activity of the crude ethanolic extract of *Brassica rapa* subsp. *chinensis* against *E. coli, S. enteritidis*, and *S. aureus*. It also seeks to identify the functional groups present in the extract using Fourier-Transform Infrared (FTIR) Spectroscopy and to compare its activity with standard antibiotics through the Kirby-Bauer disk diffusion method.

By examining the antimicrobial potential of Chinese cabbage, this research contributes to the growing body of literature on plant-derived antibacterial agents and supports ongoing efforts to develop alternative therapies for combating bacterial infections and antimicrobial resistance.

## MATERIALS AND METHOD

### Research Design

This study utilized an experimental research design to evaluate the antibacterial activity of Chinese cabbage (*Brassica rapa* subsp. *chinensis*) against selected foodborne pathogens. The methodologies employed included Fourier-Transform Infrared (FTIR) Spectroscopy for phytochemical profiling and the Kirby-Bauer disk diffusion method for antibacterial screening.

### Research Locale

The study was conducted at the University of the Immaculate Conception. Maceration and extraction of plant materials were carried out in the Analytical Chemistry Laboratory, while antimicrobial testing was performed in the Pharmaceutical Microbiology Laboratory of the said institution. FTIR analysis was conducted at the Ateneo de Davao University Chemical Laboratory. Rotary evaporation of the extract was done at the Philippine Institute of Traditional and Alternative Health Care (PITAHC).

### Collection and Preparation of Plant Material

Fresh Chinese cabbage leaves were obtained from local farmers in Marahan, Davao City. The leaves were thoroughly washed to remove dirt and contaminants, then oven-dried at 60⍰°C. A total of 500 grams of dried leaves were macerated in 80% ethanol for 24–48 hours at room temperature. The filtrate was then rotary-evaporated at 45⍰°C to concentrate the extract and lyophilized, which was stored at 4⍰°C until further use.

### Fourier-Transform Infrared Spectroscopy (FTIR)

A portion of the lyophilized ethanolic extract was subjected to FTIR spectroscopy. The procedure aimed to identify the functional groups present in the extract by analyzing characteristic absorption peaks in the infrared region.

### Test Organisms

The following bacterial strains were used in this study: *Escherichia coli* and *Salmonella enteritidis* (acquired from the Department of Science and Technology), and *Staphylococcus aureus* (acquired from the Clinical Laboratory of the University of the Immaculate Conception).

### Antibacterial Assay (Kirby-Bauer Disk Diffusion Method)

Mueller-Hinton Agar (MHA) plates were prepared and inoculated with test organisms using sterile cotton swabs. Sterile paper disks were impregnated with the plant extract and placed onto the inoculated plates. Commercial antibiotic disks—ciprofloxacin (for *E. coli* and *S. enteritidis*) and vancomycin (for *S. aureus*)—served as positive controls. Distilled water served as the negative control. Plates were incubated at 37⍰°C for 18–24 hours. Zones of inhibition were measured using a vernier caliper and compared against Clinical and Laboratory Standards Institute (CLSI) criteria (Patel, 2017).

### Sterilization and Biosafety

All glassware and culture media were sterilized by autoclaving at 121⍰°C and 15 psi for 15 minutes. Aseptic techniques were strictly followed throughout all microbiological procedures. Biosafety measures included the use of personal protective equipment, work within a biosafety cabinet with laminar air flow, and appropriate waste disposal protocols.

### Statistical Analysis

An independent two-sample t-test was performed to assess significant differences between the zones of inhibition of the plant extract and standard antibiotics for each test organism. A p-value of <0.05 was considered statistically significant.

## RESULTS

### FTIR Analysis of Plant Extract

Fourier-Transform Infrared (FTIR) Spectroscopy revealed five distinct absorption peaks in the ethanolic extract of *Brassica rapa* subsp. *chinensis*, indicating the presence of several functional groups (Table 1). These included alcohols (O–H stretch), alkenes (C=C stretch), and amines (C–N stretch), which are commonly associated with phytochemicals such as glucosinolates and their hydrolysis products.

**Table 1.** Absorption Peaks Observed Through FTIR and Possible Functional Groups.

| Peak no. | Group frequency (cm<sup>-1</sup>) | Origin | Possible Functional Group |
| --- | --- | --- | --- |
| 1 | 1066.56 | C-N | Primary/secondary/tertiary amines |
| 2 | 1396.37 | C-O | Alkoxy, alcohol |
| 3 | 1610.45 | C=C | Alkene |
| 4 | 2945.10 | C-H | Alkane, alkene, alkyl |
| 5 | 3325.05 | O-H | Alcohol |

### Antibacterial Activity (Disk Diffusion Method)

The Kirby-Bauer method was employed to evaluate the antibacterial potential of the plant extract against *E. coli, S. enteritidis*, and *S. aureus*. Zones of inhibition were measured and compared with those produced by standard antibiotics and a negative control (distilled water). Results are presented in Table 2.

**Table 2.**
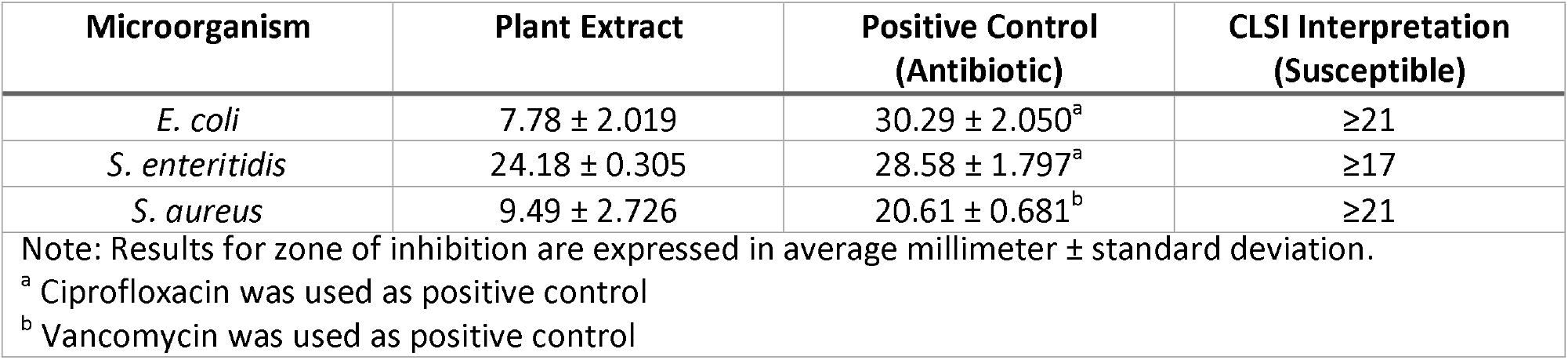
Zones of Inhibition Observed Using Kirby-Bauer Disk Diffusion Method.

**Table 3.** Statistical Report Comparing the Zones of Inhibition Between the Plant Extract and Positive Controls.

| Microorganism | t-value | p-value | Remarks |
| --- | --- | --- | --- |
| *E. coli* | 13.551 | .0002 | Significant |
| *S. enteritidis* | 4.186 | .0139 | Significant |
| *S. aureus* | 6.581 | .0024 | Significant |

Among the tested bacteria, *S. enteritidis* showed the greatest susceptibility to the plant extract, with a zone of inhibition above the CLSI threshold for susceptibility. In contrast, *E. coli* and *S. aureus* demonstrated relatively smaller zones, indicating possible resistance.

### Statistical Analysis (T-test)

Independent t-tests were conducted to assess whether the zones of inhibition produced by the plant extract were significantly different from those of standard antibiotics. All p-values obtained were less than 0.05, indicating statistically significant differences.

## DISCUSSIONS

The FTIR analysis of the Chinese cabbage (*Brassica rapa* subsp. *chinensis*) ethanolic extract revealed the presence of alcohol, alkene, amine, and alkyl functional groups. These findings are consistent with other Brassicaceae species, such as *Brassica oleracea* and *Brassica juncea*, which have shown similar spectral characteristics in previous studies (Li et al., 2013; Parikh & Khanna, 2014). The presence of these groups suggests that the extract may contain bioactive compounds such as glucosinolates and their hydrolysis products, known for their antimicrobial activity. However, FTIR alone cannot confirm the identity of specific compounds, and further phytochemical isolation would be necessary.

The antibacterial activity observed through the Kirby-Bauer disk diffusion method demonstrated that *Brassica rapa* subsp. *chinensis* possesses measurable inhibitory effects, particularly against *Salmonella enteritidis*, which exhibited a zone of inhibition above the CLSI threshold for susceptibility. This aligns with the findings of one study that reported that cabbage leaf juice exhibited inhibitory activity against *Salmonella* species (Brandi et al., 2006). In contrast, *E. coli* and *S. aureus* showed smaller inhibition zones, which may suggest moderate to low sensitivity to the extract. We postulate this may be a result of solvent choice as there is evidence to show that Chinese cabbage extracts in chloroform showed greater inhibition compared to ethanolic extracts, which produced little to no activity against *E. coli* and *S. aureus* (Rubab et al., 2018). Thus, the solvent used in this study (80% ethanol) may have influenced the observed potency.

Statistical analysis further confirmed that the antibacterial activity of the plant extract was significantly different from the positive controls (ciprofloxacin and vancomycin), indicating that while activity was present, it was not as effective as standard antibiotics. This is expected, as the plant extract is generally crude mixtures of various phytochemicals, whereas standard antibiotics are purified agents with known mechanisms of action.

Overall, the findings support the hypothesis that *Brassica rapa* subsp. *chinensis* exhibits antibacterial activity, particularly against *Salmonella enteritidis*, and contains functional groups associated with bioactive phytochemicals. These results provide a scientific basis for its traditional use and potential for development as a complementary antimicrobial agent. However, further studies involving compound isolation, quantification, and mechanism-of-action assays are necessary to fully evaluate its therapeutic potential.

## LIMITATIONS

This study was limited to in vitro evaluation of the antibacterial activity of the ethanolic extract of *Brassica rapa* subsp. *chinensis* against three selected foodborne pathogens: *Escherichia coli, Salmonella enteritidis*, and *Staphylococcus aureus*. The use of crude plant extract, rather than isolated compounds, may have affected the specificity and potency of the observed activity due to the presence of inactive or interfering constituents. Additionally, only one extraction solvent (80% ethanol) was used, which may not have been the most effective in extracting certain bioactive compounds.

The study also relied on basic phytochemical profiling through FTIR spectroscopy, which, while useful for identifying functional groups, does not provide detailed compound identification or quantification. Moreover, the absence of in vivo testing limits the ability to infer the extract’s pharmacological effects, safety, or potential toxicity in biological systems.

Lastly, the antimicrobial evaluation was confined to a limited number of bacterial strains and did not include fungal pathogens or multi-drug resistant isolates, which may be relevant for broader clinical or food safety applications.

## CONCLUSION

This study demonstrated that the ethanolic extract of Chinese cabbage (*Brassica rapa* subsp. *chinensis*) contains functional groups such as alcohols, alkenes, and amines, which are commonly associated with antimicrobial phytochemicals found in the Brassicaceae family. Through the Kirby-Bauer disk diffusion method, the extract exhibited antibacterial activity against *Salmonella enteritidis, Escherichia coli*, and *Staphylococcus aureus*, with the highest susceptibility observed in *S. enteritidis*. However, statistical analysis revealed that the plant extract was significantly less effective than standard antibiotics ciprofloxacin and vancomycin.

These findings suggest that while *Brassica rapa* subsp. *chinensis* possesses inherent antibacterial properties, its efficacy in crude form is limited when compared to conventional antimicrobial agents. Nonetheless, the presence of bioactive functional groups supports its potential as a natural source for antimicrobial compound discovery.

## RECOMMENDATIONS

In light of the study’s findings, it is recommended that future research focus on isolating and identifying the specific compounds responsible for the antibacterial activity of *Brassica rapa* subsp. *chinensis*. Alternative solvents and extraction techniques should be explored to enhance the potency of the extract.

Further studies should also investigate the mechanism of action of the active components, as well as their efficacy in in vivo models to evaluate safety and therapeutic potential. The development of topical or preservative formulations may offer practical applications, particularly in food safety or external antibacterial treatments.

## REFERENCES

Bangcongco, K. (2012, January 31). Davao City vendors insist they’re selling safe food items [News]. MindaNews. https://mindanews.com/top-stories/2012/01/davao-city-vendors-insist-theyre-selling-safe-food-items/

Brandi, G., Amagliani, G., Schiavano, G. F., De Santi, M., & Sisti, M. (2006). Activity of Brassica oleracea leaf juice on foodborne pathogenic bacteria. Journal of Food Protection, 69(9), 2274–2279. 10.4315/0362-028x-69.9.2274

Collado, L. S., Corke, H., & Dizon, E. I. (2015). Food safety in the Philippines: Problems and solutions. Quality Assurance and Safety of Crops & Foods, 45–56. 10.3920/QAS2014.x008

Havelaar, A. H., Kirk, M. D., Torgerson, P. R., Gibb, H. J., Hald, T., Lake, R. J., Praet, N., Bellinger, D. C., Silva, N. R. de, Gargouri, N., Speybroeck, N., Cawthorne, A., Mathers, C., Stein, C., Angulo, F. J., Devleesschauwer, B., & Group, on behalf of W. H. O. F. D. B. E. R. (2015). World Health Organization Global Estimates and Regional Comparisons of the Burden of Foodborne Disease in 2010. PLOS Medicine, 12(12), e1001923. 10.1371/journal.pmed.1001923

Hifnawy, M. S., Salam, R. M. A., Rabeh, M. A., & Aboseada, M. A. (2013). Glucosinolates, Glycosidically Bound Volatiles and Antimicrobial Activity of Brassica oleraceae Var. Botrytis, (Soultany Cultivar). Journal of Biology, 3(17).

Hoffmann, S., Batz, M. B., & Morris, J. G. (2012). Annual Cost of Illness and Quality-Adjusted Life Year Losses in the United States Due to 14 Foodborne Pathogens†. Journal of Food Protection, 75(7), 1292–1302. 10.4315/0362-028X.JFP-11-417

Hu, S.-H., Wang, J.-C., Kung, H.-F., Wang, J.-T., Lee, W.-L., & Yang, Y.-H. (2004). Antimicrobial effect of extracts of cruciferous vegetables. The Kaohsiung Journal of Medical Sciences, 20(12), 591–599. 10.1016/S1607-551X(09)70264-5

Johnson, I. T. (2002). Glucosinolates: Bioavailability and importance to health. International Journal for Vitamin and Nutrition Research. Internationale Zeitschrift Fur Vitamin-Und Ernahrungsforschung. Journal International De Vitaminologie Et De Nutrition, 72(1), 26–31. 10.1024/0300-9831.72.1.26

Kaur, R., Rampal, G., & Vig, A. P. (2011). Evaluation of antifungal and antioxidative potential of hydrolytic products of glucosinolates from some members of Brassicaceae family. Journal of Plant Breeding and Crop Science, 3(10), 218–228.

Kyung, K. H., & Fleming, H. P. (1997). Antimicrobial activity of sulfur compounds derived from cabbage. Journal of Food Protection, 60(1), 67–71. 10.4315/0362-028x-60.1.67

Li, B., Shan, C. L., Qiu, H., Fang, Y., Ge, M. Y., Wang, Y. L., Guo, L. B., Wu, G. X., Ibrahim, M., Xie, G. L., & Sun, G. C. (2013). Characterization of Chinese Cabbage Clubroot by Fourier Transform Infrared Spectra. Asian Journal of Chemistry, 25(15), 8460–8462. 10.14233/ajchem.2013.14790

Lubos, L. C. (2012). Microbiological Analyses of Selected Street Food Sauce Sold in Cagayan de Oro City, Philippines. IAMURE International Journal of Social Sciences, 4(1), 1–1. https://ejournals.ph/article.php?id=2237

Manguiat, L. S., & Fang, T. J. (2013). Microbiological quality of chicken- and pork-based street-vended foods from Taichung, Taiwan, and Laguna, Philippines. Food Microbiology, 36(1), 57–62. 10.1016/j.fm.2013.04.005

Pacheco-Cano, R. D., Salcedo-Hernández, R., López-Meza, J. E., Bideshi, D. K., & Barboza-Corona, J. E. (2018). Antimicrobial activity of broccoli (Brassica oleracea var. Italica) cultivar Avenger against pathogenic bacteria, phytopathogenic filamentous fungi and yeast. Journal of Applied Microbiology, 124(1), 126–135. 10.1111/jam.13629

Parikh, H., & Khanna, A. (2014). Pharmacognosy and Phytochemical Analysis of Brassica juncea Seeds. Pharmacognosy Journal, 6(5), 47–54.

Patel, J. B. (2017). Performance Standards for Antimicrobial Susceptibility Testing. Clinical and Laboratory Standards Institute.

Rubab, M., Chellia, R., Saravanakumar, K., Mandava, S., Khan, I., Tango, C. N., Hussain, M. S., Daliri, E. B.-M., Kim, S.-H., Ramakrishnan, S. R., Wang, M.-H., Lee, J., Kwon, J.-H., Chandrashekar, S., & Oh, D.-H. (2018). Preservative effect of Chinese cabbage (Brassica rapa subsp. Pekinensis) extract on their molecular docking, antioxidant and antimicrobial properties. PloS One, 13(10), e0203306. 10.1371/journal.pone.0203306

Sánchez-Pujante, P. J., Borja-Martínez, M., Pedreño, M. Á., & Almagro, L. (2017). Biosynthesis and bioactivity of glucosinolates and their production in plant in vitro cultures. Planta, 246(1), 19–32. 10.1007/s00425-017-2705-9

Scallan, E., Hoekstra, R. M., Angulo, F. J., Tauxe, R. V., Widdowson, M.-A., Roy, S. L., Jones, J. L., & Griffin, P. M. (2011). Foodborne Illness Acquired in the United States—Major Pathogens—Volume 17, Number 1—January 2011—Emerging Infectious Diseases journal—CDC. Emerging Infectious Diseases, 17(1). 10.3201/eid1701.p11101

Vale, A. P., Santos, J., Melia, N., Peixoto, V., Brito, N. V., & Oliveira, M. B. P. P. (2015). Phytochemical composition and antimicrobial properties of four varieties of Brassica oleracea sprouts. Food Control, 55, 248–256. 10.1016/j.foodcont.2015.01.051

World Health Organization. (2015). WHO estimates of the global burden of foodborne diseases: Foodborne diseases burden epidemiology reference group 2007-2015. World Health Organization. https://www.who.int/publications/i/item/9789241565165

